# Feasibility of Stereo EEG Based Brain Computer Interfacing in An Adult and Pediatric Cohort

**DOI:** 10.1101/2024.06.12.598257

**Authors:** Michael A Jensen, Gerwin Schalk, Nuri Ince, Dora Hermes, Peter Brunner, Kai J Miller

## Abstract

**Introduction:** Stereoelectroencephalography (sEEG) is a mesoscale intracranial monitoring method which records from the brain volumetrically with depth electrodes. Implementation of sEEG in BCI has not been well-described across a diverse patient cohort.

**Methods:** Across eighteen subjects, channels with high frequency broadband (HFB, 65-115Hz) power increases during hand, tongue, or foot movements during a motor screening task were provided real-time feedback based on these HFB power changes to control a cursor on a screen.

**Results:** Seventeen subjects established successful control of the overt motor BCI, but only nine were able to control imagery BCI with *≥* 80% accuracy. In successful imagery BCI, HFB power in the two target conditions separated into distinct subpopulations, which appear to engage unique subnetworks of the motor cortex compared to cued movement or imagery alone.

**Conclusion:** sEEG-based motor BCI utilizing overt movement and kinesthetic imagery is robust across patient ages and cortical regions with substantial differences in learning proficiency between real or imagined movement.

## INTRODUCTION

Brain–computer interfacing (BCI) requires a signal that is strongly correlated to a behavioral state such as movement or speech. Many types of electrical signals can be used for real-time BCI, including scalp electroencephalography (EEG)[1], magnetoencephalography (MEG)[2], electrocorticography (ECoG)[3, 4], and single neuron recordings[5, 6]. Stereoelectroencephalography (sEEG) is a mesoscale measurement that records from the brain volumetrically using depth electrodes[7]. Like ECoG, it represents an intracranial population measure of the summation of local field potentials generated from the n-poles of 100,000s of neurons surrounding the recording electrode. Compared to ECoG, sEEG is not limited to the surface of the cortex. Thus, sEEG allows for sampling from distance cortical and subcortical regions that were not previously possible with ECoG.

Currently, sEEG is utilized in the treatment of drug-resistant epilepsy. Once implantated with sEEG depth electrodes, patients remain in the hospital for characterization of their seizures. This often takes days to weeks, allowing patients to participate in experiments including brain computer interfaces, if they wish to. Historically, researchers have used spectral changes on the cortical surface to provide feedback [3, 4], allowing individuals to control a cursor on a computer screen in a matter of minutes. Our work describes the extension of this work to sEEG, including its design, implementation, and feasibility.

## MATERIALS AND METHODS

### Ethics statement

The study was approved by the Institutional Review Board of the Mayo Clinic (IRB 15-006530) and conducted according to the guidelines of the Declaration of Helsinki. Each patient or their parental guardian provided informed consent as approved by the IRB.

### Subjects

Eighteen patients (8 females, 6-37 years of age, Table 1) with drug resistant eilepsy participated in this study, after implantation with 10-17 sEEG electrode leads. Electrode planning was performed by the clinical epilepsy team using brain imaging, typical semiology, and scalp EEG. Electrode locations were not modified to accommodate research; no extra electrodes were added. All experiments were performed in the epilepsy monitoring unit (EMU) or Pediatric Intensive Care Unit (PICU) at the Mayo Clinic in Rochester, MN.

**Table 1.**
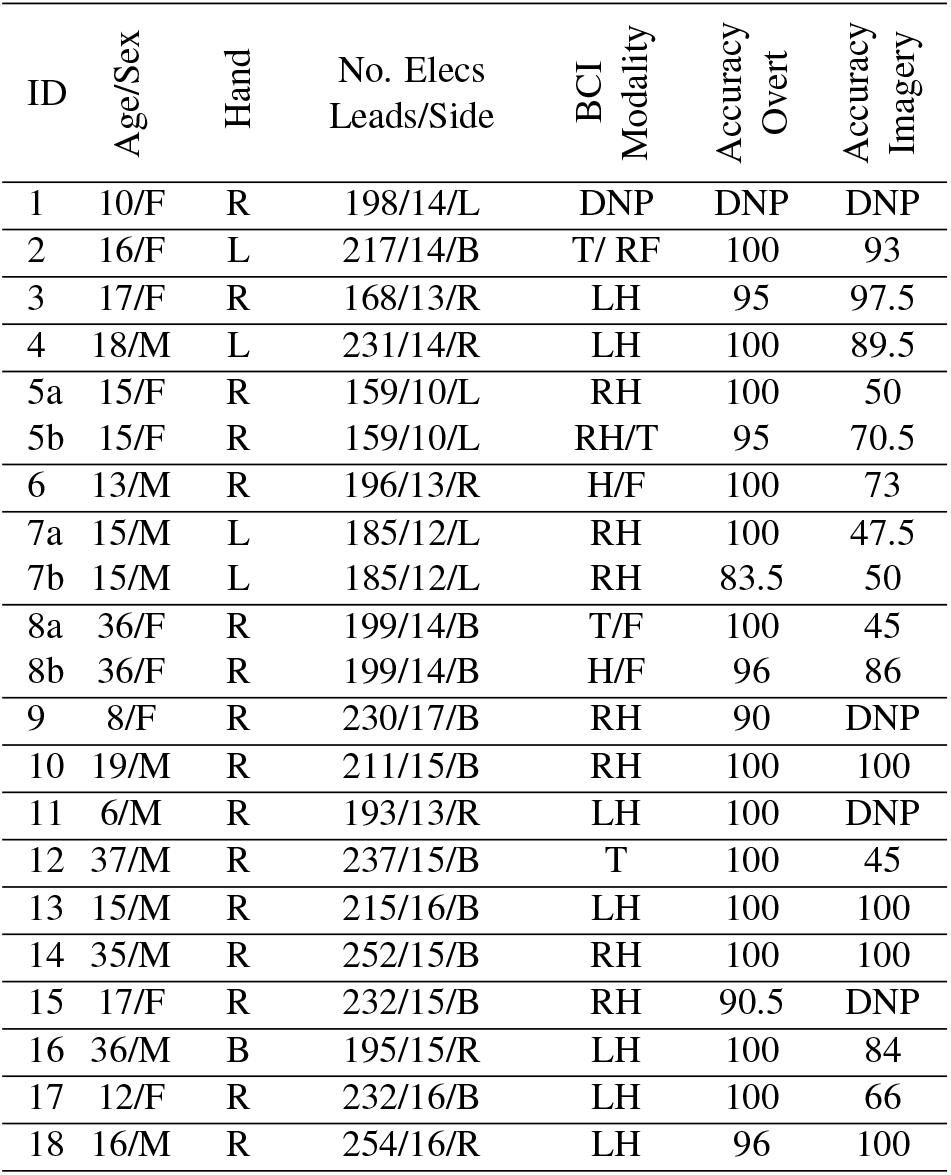
Subject Information, DNP = did not participate, “/” indicates that modalities pushed cursor in opposing directions.

| ID | Age/Sex | Hand | No. Elecs Leads/Side | BCI Modality | Accuracy Overt | Accuracy Imagery |
| --- | --- | --- | --- | --- | --- | --- |
| 1 | 10/F | R | 198/14/L | DNP | DNP | DNP |
| 2 | 16/F | L | 217/14/B | T/ RF | 100 | 93 |
| 3 | 17/F | R | 168/13/R | LH | 95 | 97.5 |
| 4 | 18/M | L | 231/14/R | LH | 100 | 89.5 |
| 5a | 15/F | R | 159/10/L | RH | 100 | 50 |
| 5b | 15/F | R | 159/10/L | RH/T | 95 | 70.5 |
| 6 | 13/M | R | 196/13/R | H/F | 100 | 73 |
| 7a | 15/M | L | 185/12/L | RH | 100 | 47.5 |
| 7b | 15/M | L | 185/12/L | RH | 83.5 | 50 |
| 8a | 36/F | R | 199/14/B | T/F | 100 | 45 |
| 8b | 36/F | R | 199/14/B | H/F | 96 | 86 |
| 9 | 8/F | R | 230/17/B | RH | 90 | DNP |
| 10 | 19/M | R | 211/15/B | RH | 100 | 100 |
| 11 | 6/M | R | 193/13/R | LH | 100 | DNP |
| 12 | 37/M | R | 237/15/B | T | 100 | 45 |
| 13 | 15/M | R | 215/16/B | LH | 100 | 100 |
| 14 | 35/M | R | 252/15/B | RH | 100 | 100 |
| 15 | 17/F | R | 232/15/B | RH | 90.5 | DNP |
| 16 | 36/M | B | 195/15/R | LH | 100 | 84 |
| 17 | 12/F | R | 232/16/B | LH | 100 | 66 |
| 18 | 16/M | R | 254/16/R | LH | 96 | 100 |

### Lead Placement, Electrode Localization, Rereferencing

Platinum depth electrode contacts (DIXI Medical) were 0.8mm in diameter with 10-18 2mm length circumferential contacts separated by 1.5mm (Fig 1). Surgical targeting and implantation were performed in the standard clinical fashion. Anatomic locations of electrodes were determined using the steps and tools described previously[8, 9]. All data were bipolar re-referenced such that channels reflect mixed activity at *two* adjacent electrode contact sites (Figs 1-4). These dipolar channels were plotted using SEEGVIEW, which slices brain renderings, and projects channels to the center of the closest slice [9] in order to present analyses in a more clinically familiar manner.

**Figure 1.**
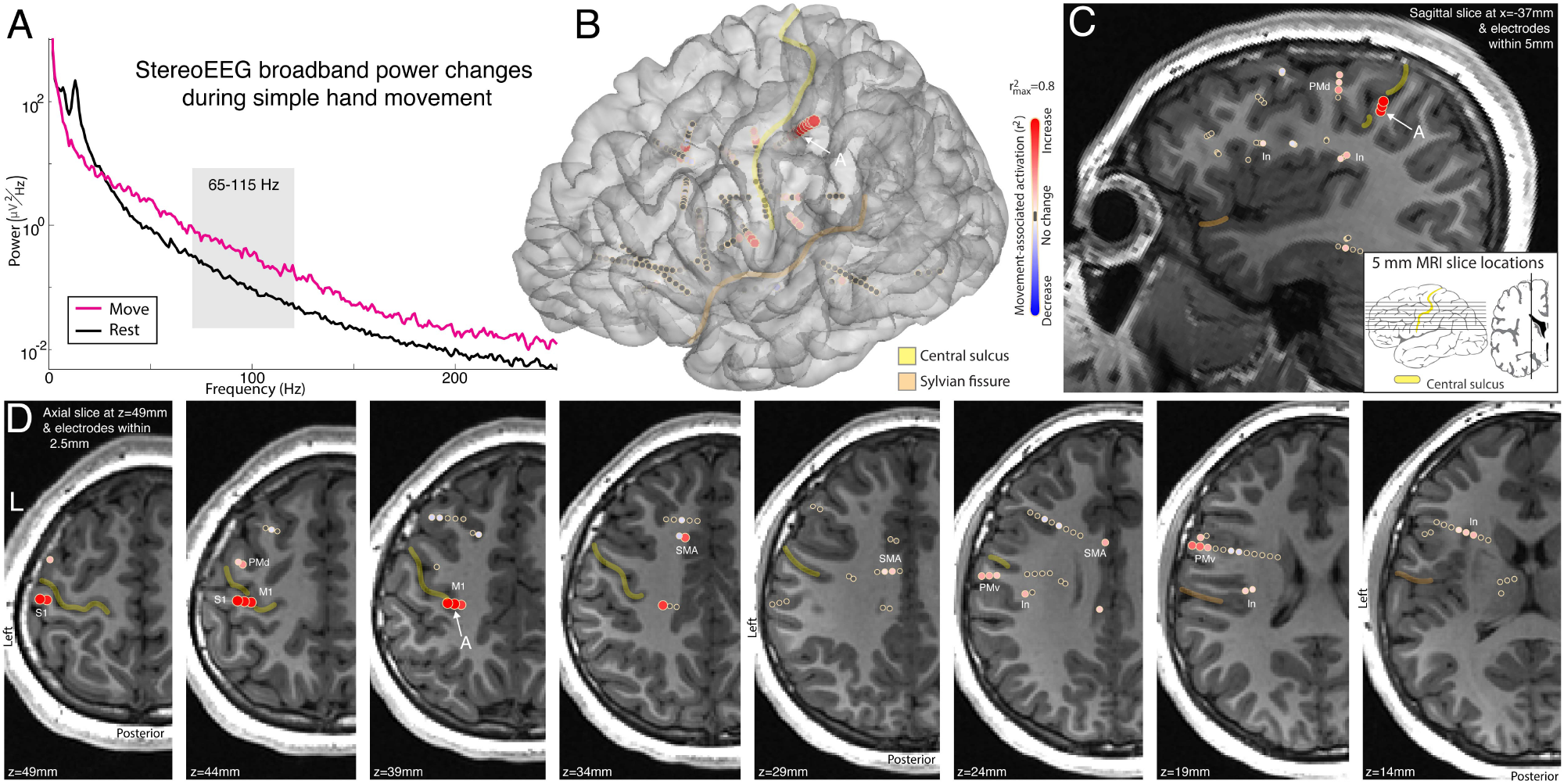
sEEG recordings during movement - Subject 1. **(A)** Power spectrum from a SEEG electrode at the sulcal base of primary motor cortex during hand movement (pink) and rest (black), from the recording site noted in panels B-D. **(B)** Broadband power (approximated by 65-115Hz band) increases during movement compared to rest. **(C)** Sagittal slice showing electrodes within 5mm of this slice allows viewing broadband power increases on the surface and at depth. **(D)** As in (C), but for axial slices and electrodes within 2.5mm. Activation maps for movement are shown in the central colorbar (signed r^2^, scaled to 1 maximum, with red/blue reflecting power increase/ decrease with movement). Yellow and peach in B-D indicate the central & sylvian fissures. Note the simultaneous measurement of M1, PMd, PMv, Insula (In), SMA, and S1 (primary sensory), which all show movement-associated broadband power increases.

### Motor Screening Task

Our motor task involved 3 seconds of 1) opening and closing of the hand, 2) side-to-side movement of the tongue with mouth closed, and 3) alternating dorsi- and plantar flexion of the foot with 3 second rest periods interleaved as described previously [8]. The BCI2000 software was used for stimulus presentation and synchronization of (EMG) and sEEG signals [10].

### Offline Signal Processing and Analysis

All analyses were performed in MATLAB. EMG signal was recorded in parallel to determine the precise timing of movement onset and offset in response to a visual cue. Within each movement trial, averaged power spectral densities (PSDs) were calculated from 1 to 300 Hz every 1 Hz using Welch’s averaged periodogram method with 1 second Hann windows to attenuate edge effects and 0.5 second overlap[11]. The averaged PSD for each movement or rest trial was normalized to the global mean across all trials. The PSDs were normalized in this way since brain signals of this type generally follow a 1/f power law and shape[12], so that lower frequency features dominate in the absence of normalization. From each of these normalized single trial PSDs, averaged power in a broadband high frequency band (65-115 Hz) was calculated for subsequent analysis, as previously described [8]. This band was chosen as it captures broadband activity above most oscillations and avoids ambient line noise at 60 Hz and 120 Hz.

For each bipolar re-referenced channel, signed *r*^2^ cross-correlation values (*r*^2^) of the mean spectra from 65-115 Hz were calculated for each movement modality. The *r*^2^ value of each channel was determined by comparing mean power spectra between rest and movement trials separately. The sign of each *r*^2^ indicates whether power is increasing or decreasing with movement, as illustrated by red and blue circles, respectively, in each figure.

### BCI Task

We implemented our BCI using the BCI2000[10] software, which applies a spectral estimator to incoming signals using an autoregressive model of the input, operating like a Fast Fourier Transform with a limited number of coefficients. A linear classifier was applied to the feature space of 70-110 Hz power in the channel(s) chosen for BCI to differentiate between movement (or imagined movement) and rest periods allowing for cursor control. During the initial experimental run, BCI2000 adapted this classifier based on the mean and variance of a data buffer (previous 30 sec of incoming data). The threshold was then set to the mean of the data buffer, and the velocity is set to the inverse square root of the variance of the data buffer. These parameters were then fixed for the remainder of the experiment to allow for online learning by each subject.

In overt BCI, patients controlled the cursor by moving in order to modulate cortical activity in the pre-selected channels, and in imagery BCI, the cursor was controlled using kinesthetic imagery alone (confirmed by EMG. Both overt and imagery BCIs in this study provided feed-back to channels that demonstrated the highest somatotopic tuning based on *r*^2^ values [8] during the motor screening task (Fig. 2).

**Figure 2.**
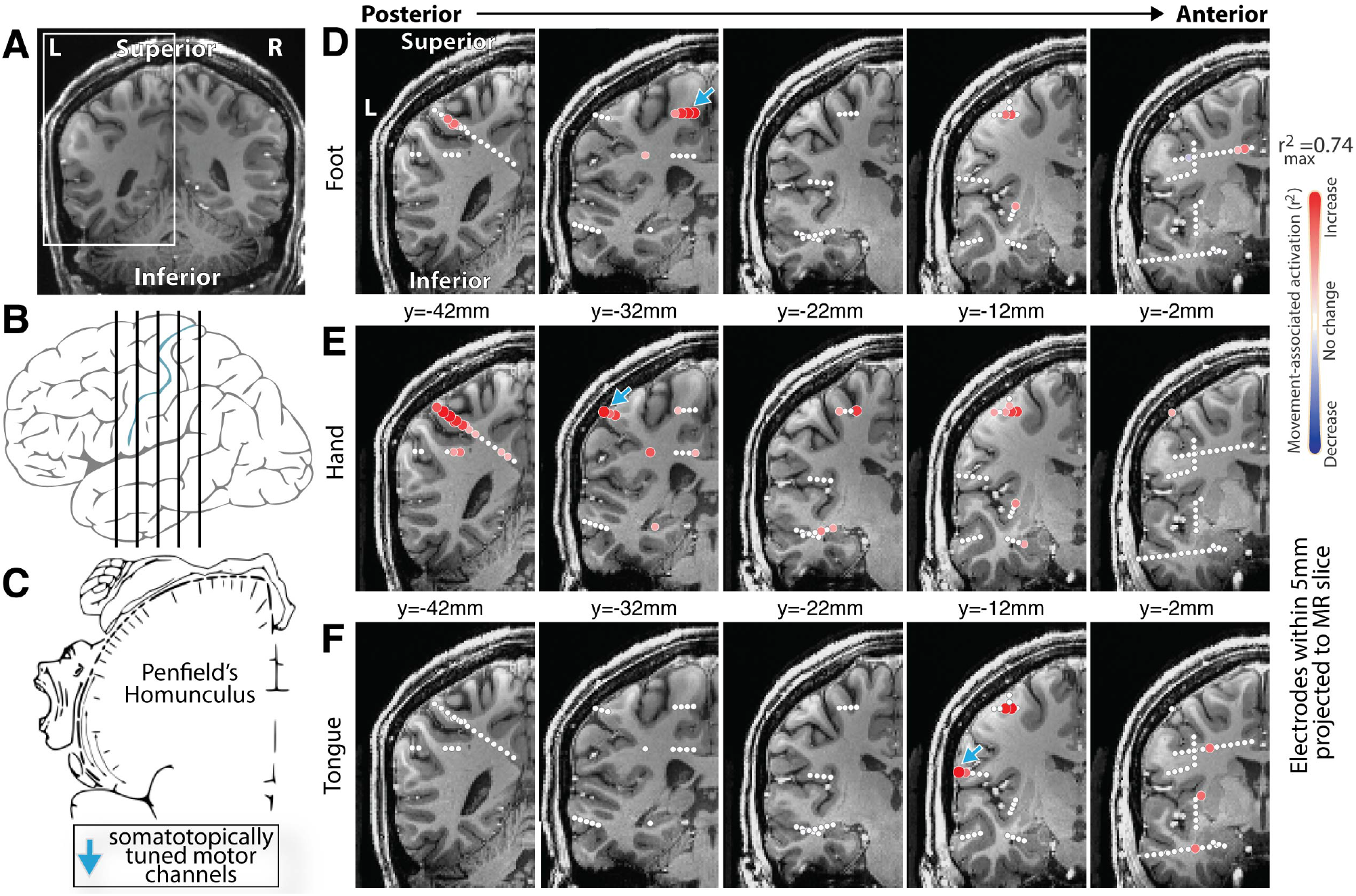
The homunculus in 3 dimensions - Subject 2. SEEG allows us to measure the volumetric structure of the homunculus electrophysiologically, shown here for the first time. **(A&B)** Locations of coronal insets in D-F. **(C)** Reproduction of Penfield’s classic motor homunculus (Wikipedia.org). **(D)** Comparison of blocks of foot movement vs rest from an SEEG array, plotting movement associated broadband (65-115Hz) change. **(E&F)** As in D, for Hand and Tongue movement. Note that the classic 2-dimensional homunculus extends into the brain depths, reflecting the volumetric nature of motor representation.

Prior to BCI, subjects were instructed to associate a target (up vs. down, left vs. right) on the screen with rest or movement (e.g. hand open/close). The target appeared at the top or bottom of the screen 1.5 seconds sec prior to a red cursor, at which point subjects proceeded to move/imagine moving or remained still once the cursor appeared (Fig. 3). Subjects were allowed 5 seconds to move the cursor to the target. If the trial was not completed (the cursor hits neither the target nor the opposite edge of the screen), this trial was not considered in the accuracy calculation and a new trial began. Each run was 2 min and allowed subjects to complete as many trials as possible. The first run was for calibration such that the computer could adapt to the power changes in the control channel(s) as subjects alternated between movements/imagined movements or rest.

**Figure 3.**
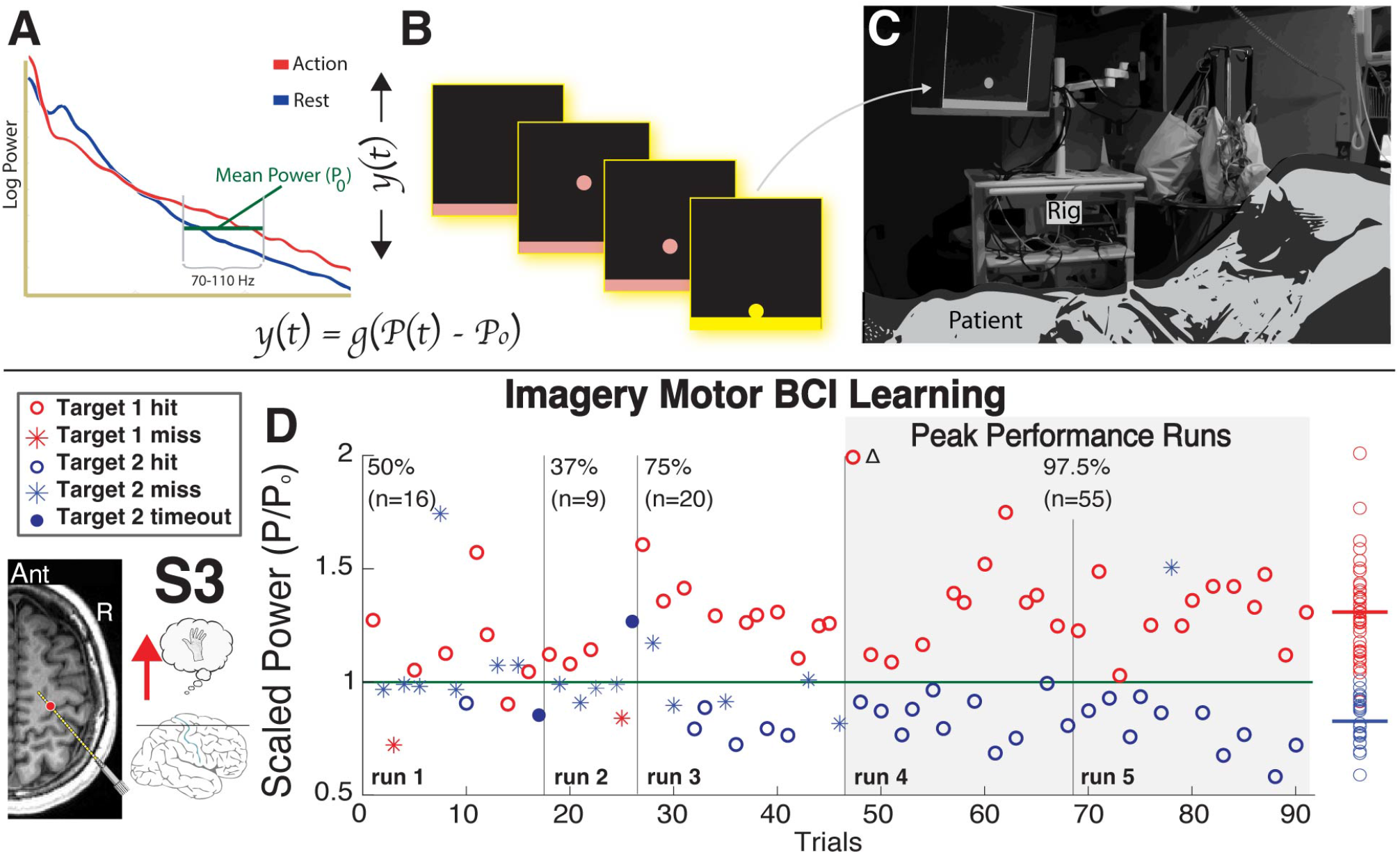
Schematic of online BCI feedback. **(A)** Power from 70-110 Hz in the channel chosen in A determines the direction and velocity of the cursor on screen. **(B)** Targets are displayed prior to cursors to cue movement or rest and subjects attempt to direct cursosrs toward the rectangular target. **(C)** Subjects perform the BCI within their bed viewing a monitor 80-100 cm from their head. **(D)** Subject 3 learned imagery BCI using a channel in the precentral gyrus across five consecutive runs. The subpopulations of power during trials of opposing targets gradually separated across the learning process until an accuracy of 97.5% was obtained (average accuracy across last two runs).

## RESULTS

### Movement

After participating in our motor screening task, changes in the power spectral density (PSD) within each sEEG channel were compared between movement and rest periods, and as in previous studies [3, 4, 8]. Movement resulted in suppression of oscillatory activity and an increase in high frequency broadband power (Fig. 1). As broadband power is correlated to local neuronal activity, it served to localize functional representation of movement across the sEEG montage (Fig 1). Germane to our goal of implementing a BCI, this enabled the identification of the somatotopically tuned cortical regions which could generate the control signal in a closed-loop feedback task (Fig. 2).

### Imagery

Subjects repeated the movement task, but were instructed to kinesthetically imagine performing the cued movement [13, 14]. As demonstrated in ECoG [3, 4], kinesthetic imagery produced an increase in broad-band power within motor regions just as during movement (Fig 5).

### BCI closed-loop feedback

Successful BCI control was defined as runs in which the cursor was moved to the correct target in *≥* 80% of trials for a minimum of 20 trials. The control channels were chosen based on the changes in HFB power associated with movement during the motor screening tasks, and it was modulation of HFB power which controlled the speed and 1-dimensional movement of a cursor on a computer monitor a few feet from the patient’s head (Fig. 3). All seventeen subjects established successful control of the overt BCI within minutes. Of these seventeen subjects, fourteen attempted to perform imagery BCI with three subjects attempting two separate BCIs for a total of seventeen. Among these subjects, nine were able to attain successful BCI control, and three controlled the cursor with above chance accuracy (Tab. 1, Fig. 4a). As represented by subject 3, learning imagery BCI occurs across several runs and results in the gradual separation of the average 70-110 Hz power within control channels between trials with opposing targets (Fig. 3). While some patients required many trials to learn the imagery BCI (Fig. 3), there was no relationship between the number of training trials and accuracy (Fig. 4b). The location of the control channel varied across patients (Tab. 1), but the majority of control channels were within the precentral gryus (PCG). Although control channels outside of the PCG may be assumed to lead to lower accuracies, control channel location did not have a strict relationship to cortical location (Fig. 4c,d).

**Figure 4.**
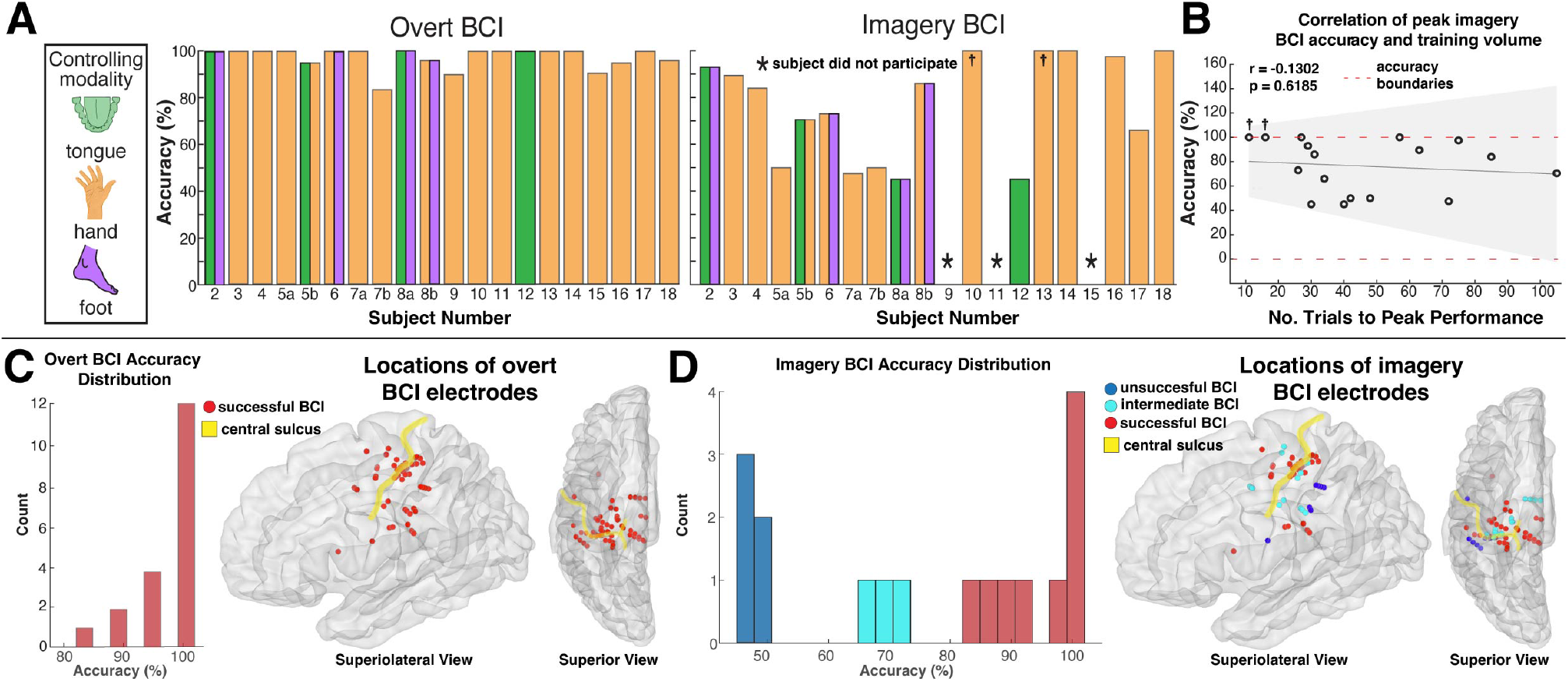
Overt and Imagery BCI accuracy across subjects. **A** Both overt (left) and imagery BCI accuracies are displayed for each subject. In some cases, subjects performed multiple BCIs that differed in either the pre-selected sEEG channels, controlling modalities, or both. Each BCI within these subjects were assigned a unique bar (e.g. subject 5a vs. 5b). † indicates subjects who attained 100% accuracy in less than 20 trials. **B** The relationship between number of training trials and peak accuracy during imagery BCI was not significant (p = 0.6185). **C** Distribution of accuracies during overt BCI (left) with locations of BCI controlling electrodes transformed to the left hemisphere of the MNI152 brain. **D** As in B, but during imagery BCI. Note that electrodes used for BCI control in subject 7 are not shown as their anatomy did not allow for accurate transformation into MNI space.

### Differential cortical engagement across tasks

Although successful BCI control necessitates broadband power modulation within the pre-selected sEEG channels controlling the BCI, activity patterns within the rest of the motor network are unconstrained. Across several subjects, we see selective engagement and differential activation based on the task being performed. For example, in Subject 3, we see maximal activity in the dorsal pre-motor area during the kinesthetic imagery screening task, and parietal engagement only when feedback in provided (Fig 5). This not only demonstrates that due to the volumetric configuration of sEEG, cortical activity across movement tasks and BCIs can be assessed on the network level, but that cortical subnetworks can be differentially engaged across tasks.

**Figure 5.**
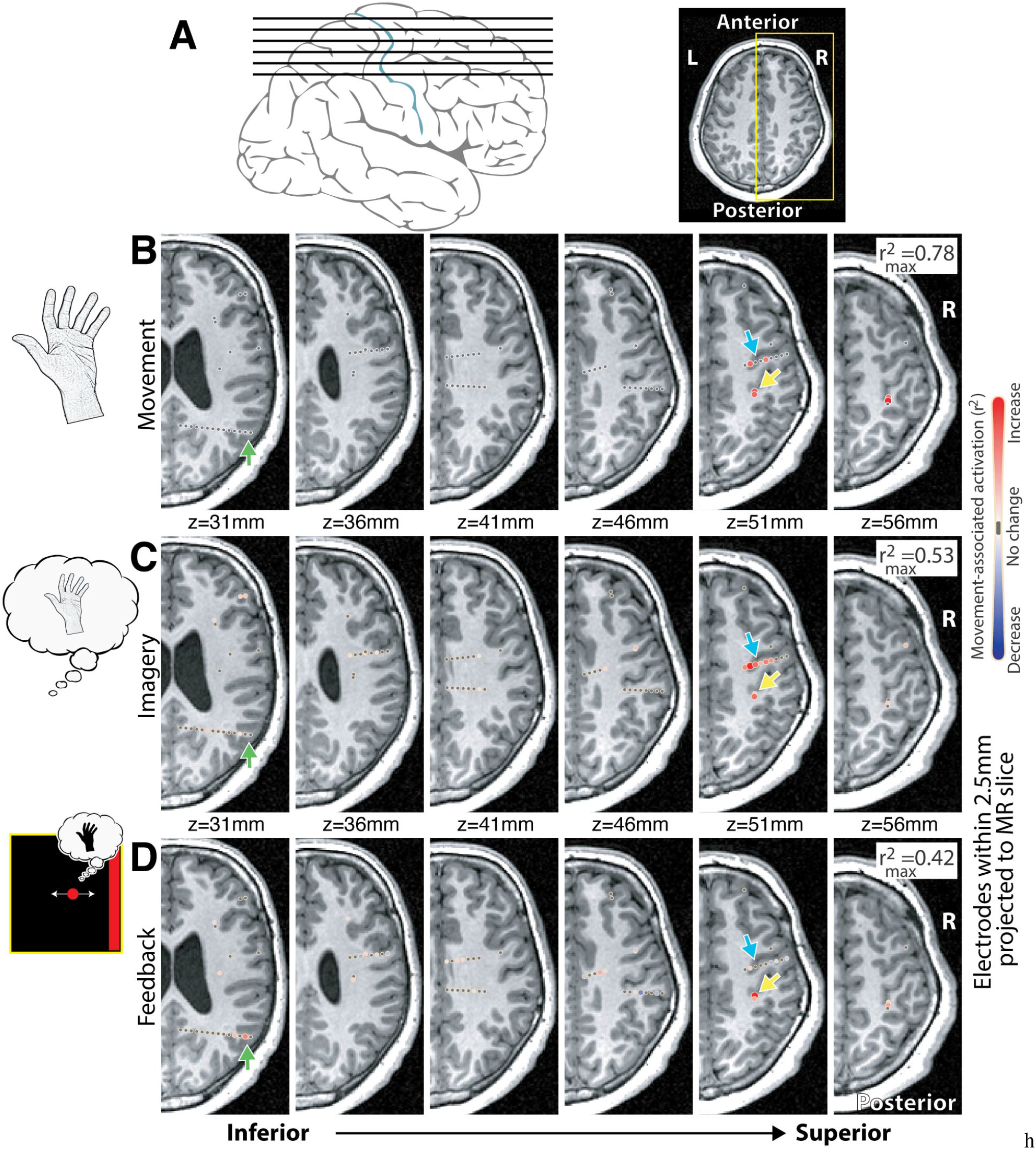
Hand movement, movement imagery, and one-dimensional BCI cursor control using sEEG - Subject 4. **(A)** Axial insets in (B-D) are as shown here. **(B)** r^2^ maps of hand movement vs rest, broadband 65-115Hz power, as in Fig 1D. **(C)** Hand movement kinesthetic imagery in the same patient. **(D)** Map of left hand imagery-based cursor control, comparing left-to-right target presentation times (cursor velocity linked linearly to 65-115Hz power from M1 site indicated by yellow arrow). Note 1) the selective augmentation in recruitment of the PMd (blue arrow) during movement imagery, and 2) the PRR (green arrow) activity selectively during BCI, but not during movement or imagery.

## DISCUSSION

Similar to ECoG studies utilizing high-frequency power to control a motor BCI[3, 4, 15], we demonstrate that both overt and imagery motor BCI can be implemented using sEEG. Even more, sEEG BCI is robust, enabling successful control of overt and imagery BCI in patients as young as six and thirteen years old, respectively. In the majority of cases in which imagery BCI accuracy was not *≥* 80%, this was due to either a lack of interest from the patient, insufficient time due the rapidly progressing clinical schedule, or an inability to learn the BCI in the allotted time.

Across the learning process, power distributions specific to time periods when each target was shown separated into two clear sub-distributions (Fig 3), with the active targets being more easily hit than inactive targets early on in learning. Presumably this may be due to the more concrete nature of kinesthetic imagery compared to rest which allows subjects to anchor to a tangible process. This is supported by results in subject 5 where the combination of imagined tongue and hand movement increased accuracy. All but two of the sEEG channels controlling the BCI were located in the pre-central gyrus (PCG). This said, selection of control channels within the PCG is not necessary nor sufficient for successful BCI control as 5 out of 8 subjects who failed to establish successful imagery BCI control used control channels in the PCG. In addition, successful imagery BCI control using channels outside of the PCG was performed in several subjects. For example, control channels for subject 4 were in the parietal operculum (Tab. 1), and control channels for subject 6 were in both primary and cingulate areas. Additionally, 14 of 17 BCI modalities involved hand movement, and of the 3 BCIs that did not utilize hand movement, 2 were unsuccessful. The disproportionate representation of the hand in our BCI is due to the sEEG trajectories chosen by the clinicians, but future work should continue to explore motor BCI modalities outside of the upper limb to allow for more powerful studies into the unique characteristics of each modalities.

Although training time was limited, there was no correlation between the number of training trials and the peak accuracy achieved by each subject (Fig 4b). This indicates that there may be a qualitative difference in the ability to learn imagery BCI across subjects independent of training volume. Certainly, the causal mechanism underlying this difference should be a explored further in future work.

Movement, kinesthetic imagery, and imagery BCI appear to differentially engage the motor network (Fig 4). While detailed exploration of this concept is outside the scope of this work, the examination of the unique roles of non-primary subnetworks of the motor network in abstract learning is a critical advantage of sEEG-based BCI.

## CONCLUSION

One-dimensional motor, sEEG-based BCI utilizing overt movement and kinesthetic imagery is robust across patient ages and cortical regions. Subjects differ in their ability to learn imagery BCI, and further work should explore the mechanism behind this difference.

## ACKNOWLEDGEMENTS

We extend our deepest gratitude to the patients who volunteered their time in the EMU to participate in our research, to Bambi Wessel, Cindy Nelson, and the staff at St. Mary’s hospital. This work was supported by the NIH U01-NS128612 (KJM, PB), Brain and Behavior Research Foundation (KJM), and the Foundation for OCD Research (KJM). The contents of this manuscript are solely the responsibility of the authors and do not necessarily represent the official views of the NIH. Our funders played no role in data collection and analysis, study design, decision to publish, or manuscript preparation.

## DATA AND CODE AVAILABILITY

At the time of final publication, all data will be made available at https://osf.io/kxqvw/. All code will be made available at the author’s GitHub https://github.com/michaeljensen42.

